# Hypoxia reduces induced LRIG1 protein despite sustained mRNA induction in colorectal cancer organoids

**DOI:** 10.64898/2026.10.19.760078

**Authors:** Akari Minamiura, Shuto Aoki, Rio Matsuwaka, Arisa Yoshimoto, Yumi Nobori, Takahiro Matsuura, Shinji Takamatsu, Eiji Miyoshi, Jumpei Kondo

**Affiliations:** Department of Molecular Biochemistry and Clinical Investigation, Division of Health Sciences, The University of Osaka, Graduate School of Medicine, Suita, Japan

**Author notes:** Corresponding Author: Jumpei Kondo Department of Molecular Biochemistry and Clinical Investigation, Division of Health Sciences, The University of Osaka, Graduate School of Medicine 1-7 Yamada-oka, Suita, Osaka 565-0871, Japan. Current Affiliation of E Miyoshi: Graduate School of Medical Safety Management, Jikei University of Health Care Sciences, Osaka, Osaka, Japan.

## Abstract

LRIG1 is a negative regulator of receptor tyrosine kinase (RTK) signaling that promotes tumor suppression by driving degradation of EGFR and other RTKs. Inducing LRIG1 expression has therefore been proposed as a strategy to limit cancer cell survival under hypoxia. However, whether hypoxia itself affects LRIG1 expression remains unclear. Here, we investigated how hypoxia affects LRIG1 expression and protein abundance in human cancer datasets and colorectal cancer organoids. Analysis of The Cancer Genome Atlas data across gastrointestinal and other cancers revealed an inverse association between *LRIG1* mRNA expression and a hypoxia gene signature that persisted after adjustment for global gene expression. We previously showed that BMPR inhibition induces LRIG1 and suppresses colorectal cancer organoid growth through LRIG1. Under hypoxia, this growth inhibition in colorectal cancer organoids was attenuated, and LRIG1 protein accumulation was reduced despite sustained mRNA induction. Pre-induced LRIG1 protein declined within 3 h of hypoxic exposure in organoids, and ectopically expressed LRIG1 similarly decreased in HEK293 cells. Pharmacological HIF1 inhibition did not prevent LRIG1 protein loss under hypoxia, whereas HIF1α stabilization did not reproduce it under normoxia, suggesting limited HIF1 involvement. Bafilomycin A1 restored LRIG1 protein levels, whereas chloroquine and the proteasome inhibitor MG132 did not, implicating a bafilomycin A1-sensitive lysosomal process. Notably, the hypoxia-induced decrease in LRIG1 was not accompanied by a decrease in total EGFR. These findings suggest that hypoxia limits LRIG1 protein induction, potentially constraining strategies that rely on LRIG1 induction in cancer treatment.

## 1. Introduction

Hypoxia is a defining feature of the tumor microenvironment, arising from rapid tumor expansion and the formation of structurally and functionally aberrant tumor vasculature[1]. The hypoxic response reprograms cancer cells in various aspects, including modification of metabolism, promotion of invasion and metastasis, and resistance to therapy[2,3]. Hypoxia-inducible factor 1 (HIF1) mediates many of these responses[2,3]: under normoxia, prolyl hydroxylase domain proteins (PHDs) target HIF1α for proteasomal degradation, whereas hypoxia inhibits PHD activity, stabilizing HIF1α and inducing HIF1-dependent gene expression[4-7].

Receptor tyrosine kinase (RTK)–driven proliferative signaling is central to cancer growth. Leucine-rich repeats and immunoglobulin-like domains 1 (LRIG1) is a transmembrane negative regulator of RTKs, including EGFR, that promotes receptor degradation and limits proliferative signaling [8-10]. We previously reported that BMP pathway inhibition induces LRIG1 expression and thereby suppresses the growth of a subset of colorectal cancer organoids through LRIG1-mediated EGFR downregulation. [11]. Regarding the significance of LRIG1 under hypoxia, studies using glioma[12] and melanoma[13] cell lines *in vitro* have shown that forced LRIG1 overexpression exerts tumor-suppressive effects under hypoxic conditions, notably by attenuating hypoxia-induced epithelial–mesenchymal transition and vasculogenic mimicry formation. These studies, however, addressed the functional consequences of LRIG1 expression rather than the mechanisms controlling its abundance, and were conducted exclusively in established cell lines. Consequently, how hypoxia regulates endogenous LRIG1 expression remains poorly understood. A conference abstract has suggested that hypoxia suppresses LRIG1 protein synthesis in a HIF1-dependent manner[14]. However, how hypoxia affects endogenous and induced LRIG1 mRNA and protein levels in colorectal cancer organoids remains unknown.

In this study, we combined analyses of clinical cancer datasets with colorectal cancer organoid models to examine how hypoxia affects LRIG1 mRNA and protein levels and to investigate the processes underlying their discordance.

## 2. Materials and Methods

### 2.1. Organoid and cell line culture

Patient-derived colorectal cancer organoids (C45 and CB3 [15]), provided by Dr. Masahiro Inoue (Kyoto University), were used with approval from the Institutional Ethics Committee of the University of Osaka Hospital (No. 21129) and in accordance with the Declaration of Helsinki. Organoids were maintained in suspension in StemPro medium (Thermo Fisher Scientific) and DMEM/Ham’s F-12 (FUJIFILM Wako) at 1:7, supplemented with 1% L-alanyl-L-glutamine (FUJIFILM Wako) and 1% penicillin-streptomycin (Nacalai Tesque). For experiments, organoids were cultured in experimental medium: Advanced DMEM/F12 (Gibco) containing the same supplements, with or without LDN-193189 (LDN, Selleck). HEK293 cells (ATCC) were cultured in DMEM high-glucose (Nacalai Tesque) with 10% FBS (Corning) and 1% penicillin-streptomycin. HEK293 cells with doxycycline-inducible expression of C-terminally FLAG-tagged LRIG1 were generated by lentiviral transduction with the previously reported LRIG1 construct [16] subcloned into pLIX_403 (a gift from Dr. David Root; Addgene plasmid #41395).

For hypoxia, organoids and HEK293 cells were cultured at 1% O_2_ in a multi-gas incubator, and normoxic controls were kept at 21% O₂ in a standard CO_2_ incubator. Inhibitors (YC-1 (Selleck), MK-8617 (Selleck), MG132 (FUJIFILM Wako), bafilomycin A1 (CST), or chloroquine (Tokyo Chemical Industry) were added 1 h before hypoxic exposure.

### 2.2. Proliferation assay

Two to three days after passaging by mechanical disruption through a 27G needle, organoids of 70–100 μm were isolated by sequential filtration through 100 μm and 70 μm cell strainers (AS ONE) and recovered from the 70 μm strainer. Organoids were seeded at 20 organoids per well (n = 6 per condition) in 96-well non-treated plates in 150 μL of experimental medium with 2% Matrigel (Corning) and cultured for 5 days. Organoids in each well were counted under a microscope immediately after seeding. Where indicated, LDN was added at 100 nM. Viability was measured with CellTiter-Glo Luminescent Cell Viability Assay (Promega) on an Infinite M Plex microplate reader (Tecan), and luminescence was normalized to the initial organoid count of each well.

### 2.3. Western blotting

Organoids were lysed in RIPA buffer containing protease and phosphatase inhibitor cocktails (Nacalai Tesque) and sonicated. Protein concentration was determined using the BCA Protein Assay (Nacalai Tesque). Lysates were boiled in 4× SDS sample buffer containing 2-mercaptoethanol and separated by SDS-PAGE (7.5% or 10% gels). Proteins were transferred to Immobilon-P PVDF membranes (0.45 μm; Millipore) by semi-dry blotting, blocked with Bullet Blocking One (Nacalai Tesque), and incubated with primary antibodies (4°C, overnight) followed by HRP-conjugated secondary antibodies (room temperature, 1 h) (Table S1). Signals were detected using Chemi-Lumi One Super (Nacalai Tesque) and a FUSION imaging system (Vilber-Lourmat).

### 2.4. Semi-quantitative reverse transcription PCR

Total RNA was extracted using the RNeasy Mini Kit (QIAGEN) according to the manufacturer’s protocol, and RNA concentration and purity were assessed with a NanoDrop spectrophotometer (Thermo Fisher Scientific). RNA was reverse-transcribed using the GoScript Reverse Transcription Mix with Oligo(dT) primers (Promega), and the resulting cDNA was used for downstream analysis. Real-time qPCR was performed using THUNDERBIRD Next SYBR qPCR Mix (Toyobo) on an Mx3000P system (Agilent Technologies), with LRIG1-specific primers (forward: 5’-TGTGTCCAAGAGATGCAAGC-3’; reverse: 5’-GCTTGTCCCCTGGAGTAACA-3’). Statistical analysis was performed with Student’s t-test with Bonferroni correction.

### 2.5. Database analysis

TCGA RNA-seq study IDs are listed in Table S2. Batch-normalized RSEM values (Illumina HiSeq_RNASeqV2) were log2-transformed for cross-cancer comparisons and visualization. To evaluate the association between LRIG1 expression and hypoxia-related transcription across cancer types, RSEM Z-scores were used. Where multiple entries shared a gene symbol, their Z-scores were averaged for each sample. The hypoxia score was defined as the mean Z-score of the genes in the Buffa hypoxia signature[17] (Table S3), and LRIG1 expression as the RSEM Z-score of LRIG1. Their associations were assessed by Pearson’s correlation. To account for potential bias attributable to overall transcriptional activity, a global expression score (the mean Z-score of all genes per sample) was also calculated. P values were adjusted for multiple comparisons using the Holm method. All analyses were performed in R (v4.5.2) using the dplyr, tidyr, readr, stringr, ggplot2, ggpubr, and reshape2 packages.

### 2.6. RNA sequencing

RNA sequencing was performed by the Genome Information Research Center, Research Institute for Microbial Diseases/Immunology Frontier Research Center, Osaka University. Libraries were prepared with the TruSeq Stranded mRNA Library Prep Kit (Illumina) and sequenced on a NovaSeq 6000 (Illumina) in 101-bp single-read mode. Reads were trimmed with Trimmomatic (v0.38), aligned with TopHat (v2.1.1), and quantified as FPKM with Cuffdiff (v2.2.1).

FPKM values were log2-transformed after adding a pseudocount of 1. For each gene set (Table S3), genes present in the expression matrix were retained and Z-scored across samples for each gene. Gene set scores were calculated as the mean Z-score of the retained genes for each sample. Lysosomal degradation, autophagy, and proteasomal degradation gene sets were visualized as heatmaps using heatmap3 in R (v4.5.2). The RNA-seq data were deposited in the NCBI Sequence Read Archive (SRA) under BioProject accession number PRJNA1537312.

## 3. Results

### 3.1. Hypoxia suppresses *LRIG1* expression across cancer types

To determine whether hypoxia influences *LRIG1* expression across tumor types, we analyzed The Cancer Genome Atlas (TCGA) RNA-seq data from gastrointestinal cancers (ESCA, STAD, COADREAD, PAAD, LIHC, and CHOL); LUAD, in which high *LRIG1* expression correlates with favorable prognosis[18]; PRAD[19,20], BRCA[21,22] and HNSC[23], in which *LRIG1* act as a tumor suppressor; and glioma[12] (GBM, LGG) and melanoma[13] (SKCM), in which hypoxia regulates *LRIG1* expression. *LRIG1* expression varied widely among tumors in all these cancer types (Fig. 1A). Hypoxia-score, derived from the expression pattern of hypoxia-associated genes, showed weak-to-moderate negative correlation with *LRIG1* expression in most cancer types (Fig. 1B, Table 1). Because hypoxia can globally suppress gene expression, we also assessed its correlation with global gene expression: with the exception of LIHC (moderate positive correlation), most cancer types showed no or only a weak negative correlation between hypoxia-score and overall gene expression (Table S4). Nevertheless, after controlling for the effect of global expression by partial correlation, the weak-to-moderate correlation between *LRIG1* and hypoxia-score persisted across all cancer types and was significant in all cancers except CHOL, which had a small sample size (Table 2). These findings indicate that *LRIG1* expression is suppressed under hypoxia in various human cancers, suggesting a proliferative advantage.

**Figure 1.**
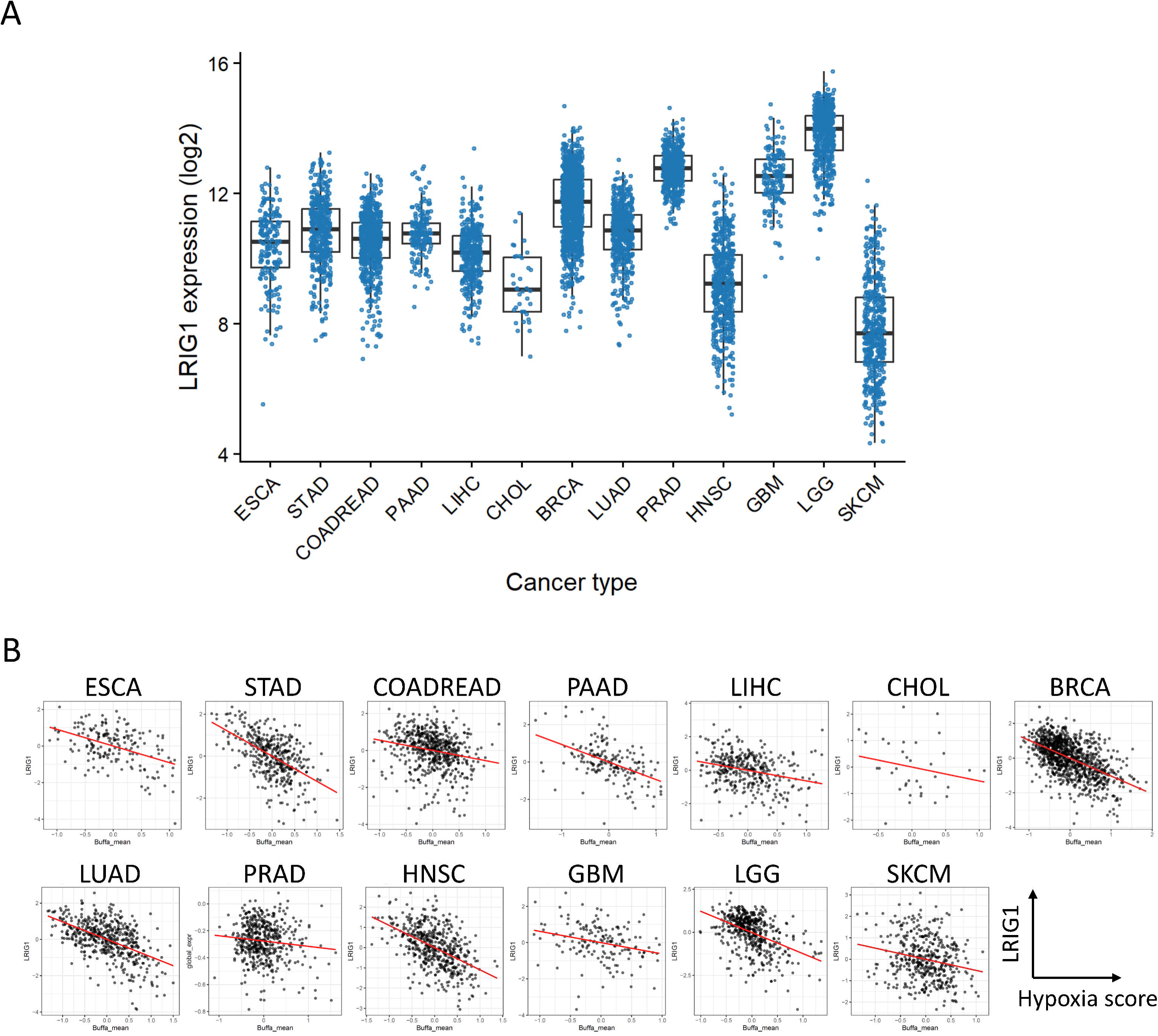
Hypoxia suppresses LRIG1 expression across cancer types (A) Pan-cancer LRIG1 gene expression. The y-axis shows mRNA expression [log2(RSEM-normalized count)]. (B) Correlation plots between LRIG1 gene expression and hypoxia score across cancer types. Red lines indicate regression curves. Sample numbers, correlation coefficients, and p-values are shown in Table S4.

### 3.2. Hypoxia uncouples LRIG1 protein from mRNA induction in colorectal cancer organoids

Although LRIG1 induction has been proposed to limit cancer cell survival under hypoxia[12,13], our TCGA analysis showed an inverse association between hypoxia and *LRIG*1 mRNA expression. We previously reported that BMP inhibition induces LRIG1 gene and protein expression in colorectal cancer (CRC) organoids under normoxia[11]. Using this model, we found that hypoxia attenuated the growth-inhibitory effect of the BMP inhibitor LDN193189 (LDN; Fig. 2A, B) and suppressed LDN-induced LRIG1 protein expression (Fig. 2C). In contrast, LDN-induced *LRIG1* mRNA expression remained robust in two representative CRC organoids under hypoxia, with no significant statistical difference from normoxia (Fig. 2D). These findings suggest that hypoxia limits LRIG1 protein induction, at least in part, through post-transcriptional regulation.

**Figure 2.**
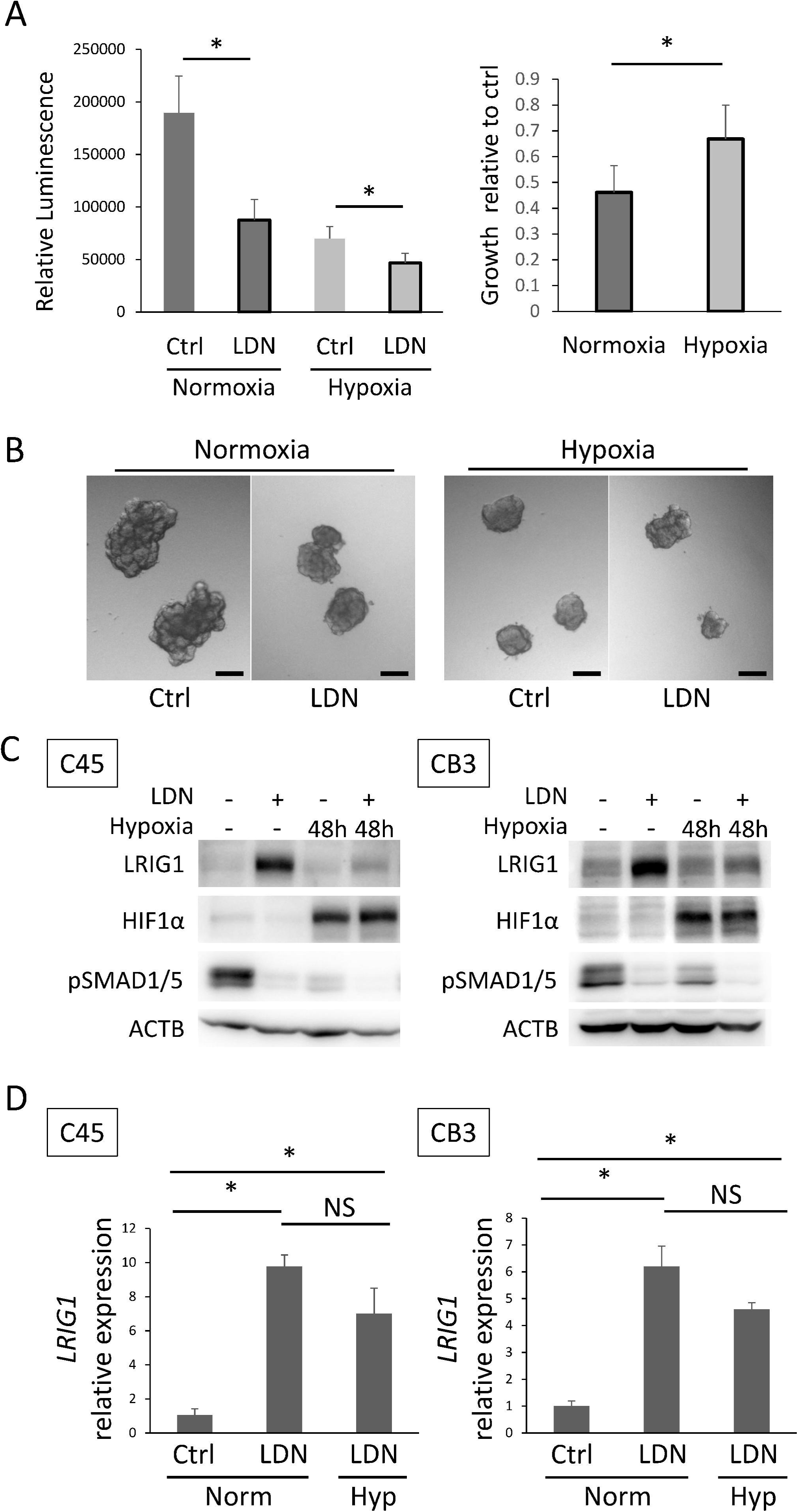
Hypoxia uncouples LRIG1 protein from mRNA induction in colorectal cancer organoids (A) ATP assay results in the colorectal cancer organoid C45 cultured under normoxia or hypoxia (1% O_2_), in the absence (ctrl) or presence (LDN) of LDN193189 treatment. Left, luminescence values normalized to organoid number at day 0. Right, organoid growth relative to the control (w/o LDN) condition at each oxygen concentration. *p < 0.05, t-test (with Bonferroni correction for multiple comparisons). (B) Representative microscopic images of the cultures shown in (A) at day 5. Scale bar, 50 μm. (C) Western blot analysis of C45 and CB3 organoids cultured for 48 hours under normoxia or hypoxia, with or without LDN treatment. (D) qPCR analysis of *LRIG1* mRNA expression in C45 and CB3 organoids. *p < 0.05, NS; not significant, t-test with Bonferroni correction.

### 3.3. Hypoxia rapidly reduces induced LRIG1 protein independently of HIF1

The reduced efficacy of BMP inhibition under hypoxia could limit its potential as a strategy to induce LRIG1 in cancer cells. To investigate the discordance between LRIG1 mRNA and protein levels, we examined how hypoxia affects LRIG1 protein that had already been induced. In the experiments shown in Fig. 2, hypoxia and BMP inhibition were initiated simultaneously, and LRIG1 protein levels were assessed 48 hours later. Here, we first induced LRIG1 under normoxia and then exposed the cells to hypoxia. LRIG1 protein levels decreased within 3 hours of hypoxic exposure in CRC organoids (Fig. 3A). Similarly, forced-expressed LRIG1 decreased rapidly in HEK293 cells (Fig. 3B), indicating that this response is not limited to BMP inhibition-induced LRIG1.

**Figure 3.**
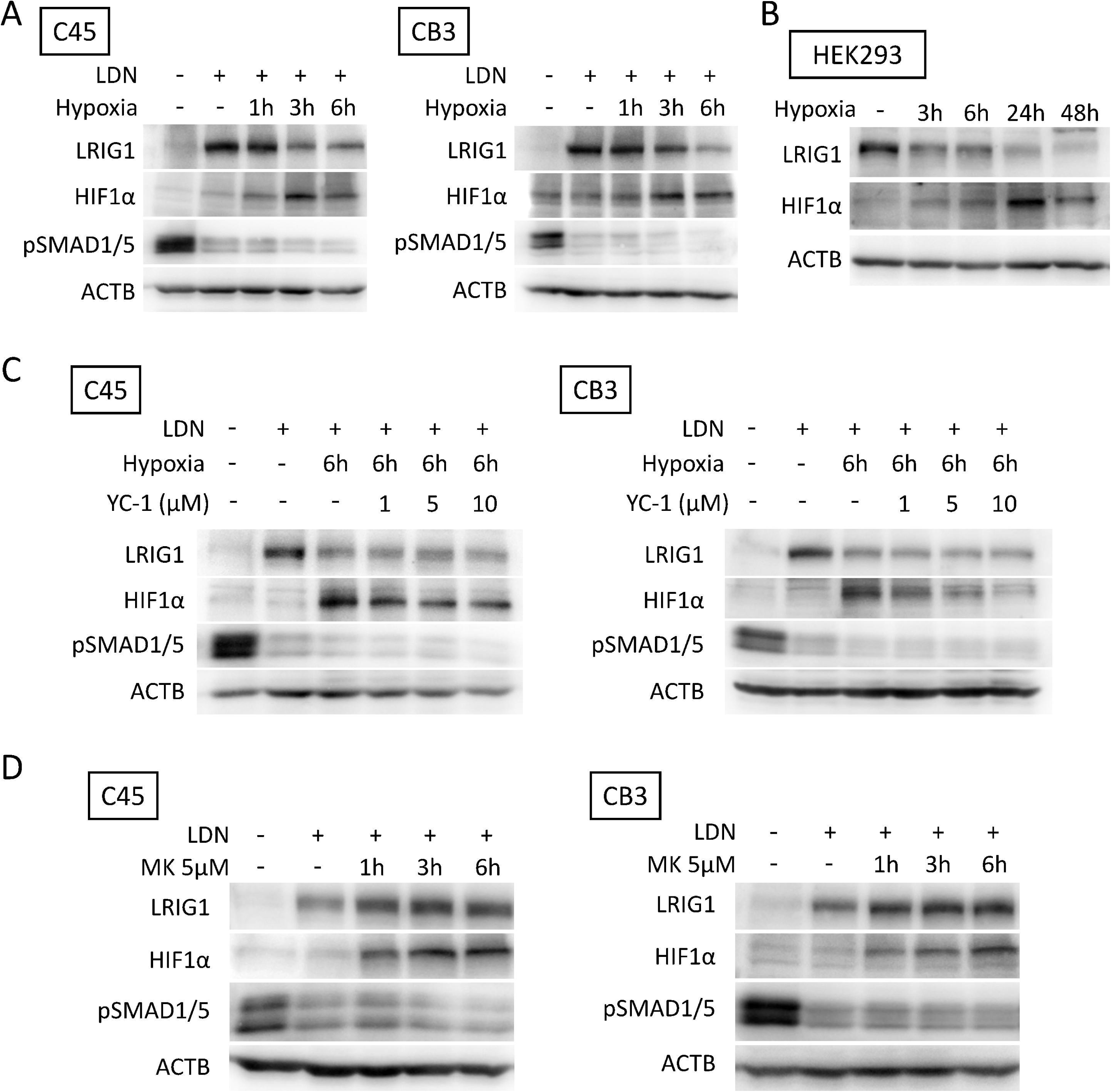
Hypoxia rapidly reduces induced LRIG1 protein independently of HIF1 Western blot analyses were performed for all panels. In (A) and (C), where indicated, LDN was added 2 days before the start of hypoxic culture and maintained throughout the exposure. (A) C45 and CB3 organoids cultured under normoxia or hypoxia. (B) HEK293 cells expressing exogenous LRIG1 cultured under normoxia or hypoxia. (C) C45 and CB3 organoids cultured under normoxia or hypoxia with or without YC-1. (D) C45 and CB3 organoids cultured under normoxia with or without MK-8617.

We next evaluated whether HIF1 is involved in the hypoxia-induced decrease in LRIG1 protein. The HIF1 inhibitor YC-1 suppressed HIF1α accumulation under hypoxia but did not prevent the decrease in LRIG1 protein (Fig. 3C). Conversely, the PHD inhibitor MK-8617 stabilized HIF1α under normoxia but did not reproduce the hypoxia-induced decrease in LRIG1 protein (Fig. 3D). Together, these findings suggest that hypoxia decreases LRIG1 protein levels independently of HIF1 activation.

### 3.4. Hypoxia-associated LRIG1 protein loss is sensitive to lysosomal inhibition

We next investigated whether the loss of LRIG1 protein under hypoxia involves enhanced degradation. To identify candidate pathways involved in LRIG1 protein loss, we analyzed the expression of proteasomal and lysosomal/autophagy-related genes in RNA-seq data from C45 CRC organoids cultured at 20% or 1% O₂ for 48 hours. The RNA-seq data confirmed both a hypoxic transcriptional response (Fig. 4A, B) and sustained LDN-induced *LRIG1* mRNA expression under hypoxia (Fig. 4C). We calculated a score for each pathway by averaging the Z-scores of its constituent genes in each sample. Under hypoxia, lysosomal and autophagy-related scores increased, whereas the proteasomal score did not (Fig. 4D). We then tested whether inhibiting these degradation pathways could preserve LRIG1 protein under hypoxia. The proteasome inhibitor MG132 did not restore LRIG1 protein levels, although it increased HIF1α accumulation, consistent with drug activity (Fig. 4E). We next used bafilomycin A1, which inhibits vacuolar H⁺-ATPase and lysosomal acidification, and chloroquine, which impairs late-stage autophagy. Bafilomycin A1 restored LRIG1 protein levels, whereas chloroquine did not, despite inducing LC3-II accumulation (Fig. 4F). Together, these findings implicate a bafilomycin A1-sensitive lysosomal process in the hypoxia-induced loss of LRIG1 protein. To further assess LRIG1 protein turnover, we performed cycloheximide-chase experiments in HEK293 cells expressing doxycycline-inducible LRIG1. LRIG1 protein declined rapidly even under normoxia and was significantly lower under hypoxia at 3 h (Fig. S1A, B). Because LRIG1 can be degraded together with EGFR during receptor downregulation[8], we asked whether the hypoxia-induced loss of LRIG1 was accompanied by EGFR loss. After 6 h of hypoxia, total EGFR levels were not reduced despite LRIG1 loss, and bafilomycin A1 did not increase EGFR levels (Fig. S1C). Together, these findings suggest that LRIG1 loss under hypoxia does not necessarily reflect concomitant EGFR loss.

**Figure 4.**
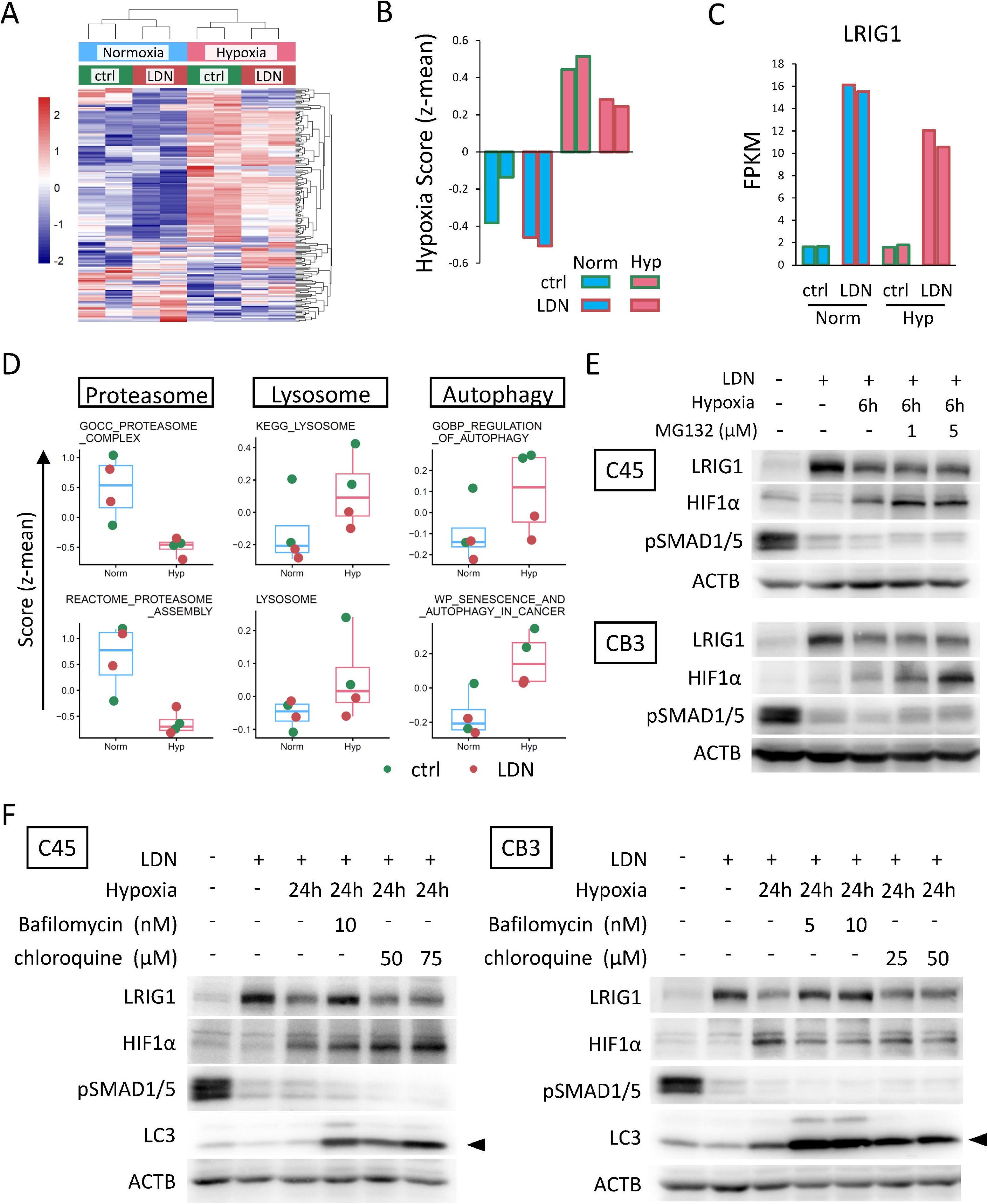
Hypoxia-associated LRIG1 protein loss is sensitive to lysosomal inhibition (A) Heatmap of differentially expressed genes in C45 colorectal cancer organoids cultured under normoxia or hypoxia with or without LDN, based on RNA-seq data. (B-C) Hypoxia scores (B) and *LRIG1* mRNA expression (C) derived from the RNA-seq data shown in (A). (D) Z-mean scores for proteasome-, lysosome-, and autophagy-related gene sets in the same RNA-seq dataset. (E-F) Western blot analysis of C45 and CB3 organoids cultured under normoxia and hypoxia with or without MG132 (E), bafilomycin A1, or chloroquine (F). Arrowheads in (F) indicate the LC3-II band size.

## 4. Discussion

LRIG1 is a negative regulator of cancer cell survival that promotes the degradation of various receptor tyrosine kinases (RTKs), including EGFR [8-11,16,21], and forced expression of LRIG1 under hypoxia has been reported to suppress tumor survival and proliferation[12,13]. In TCGA datasets, we found an inverse correlation between *LRIG1* expression and a hypoxia gene signature across multiple cancer types. In patient-derived colorectal cancer organoids, hypoxia attenuated the growth inhibition by a BMPR inhibitor LDN193189 and reduced LRIG1 protein levels. Notably, LRIG1 protein also declined rapidly during cycloheximide chase under normoxia, indicating substantial turnover even without hypoxic stress. Hypoxia further accelerated this decline, suggesting that hypoxia shifts the balance between LRIG1 protein synthesis and loss, although the relative contribution of each remains unclear. These findings together indicate that hypoxia may limit the effectiveness of approaches that rely on LRIG1 induction.

Pharmacological inhibition of HIF1 did not prevent hypoxia-associated LRIG1 loss, whereas stabilization of HIF1α under normoxia did not reproduce it. These results suggest that HIF1 activation is neither necessary nor sufficient for the observed response under the conditions tested. Hypoxia can affect protein abundance through HIF-independent pathways [24-26], but the mechanism responsible for LRIG1 loss remains to be determined. Bafilomycin A1 restored LRIG1 protein levels under hypoxia, whereas chloroquine and MG132 did not, supporting the involvement of a bafilomycin A1-sensitive lysosomal process. Endosomal trafficking followed by lysosomal degradation is a plausible route for LRIG1, as it is for other membrane proteins[27-29]. Hypoxia can alter membrane-protein trafficking, but its effects differ among proteins. Hypoxia suppresses bulk constitutive endocytosis through a HIF-independent mechanism[30], while HIF-dependent changes in endosomal trafficking can prolong RTK stability[31]. In contrast, hypoxia promotes neuropilin-1 loss through a process sensitive to both bafilomycin A1 and chloroquine[32]. The different responses of LRIG1 and neuropilin-1 to chloroquine suggest that their hypoxia-associated loss may involve different steps in endolysosomal trafficking. Determining whether hypoxia alters LRIG1 internalization, sorting, or degradation will require further analysis.

Hypoxia can enhance RTK signaling by increasing receptor expression or stability in various cancers[30,33]. In our organoids, however, hypoxia reduced LRIG1 protein levels without a corresponding decrease in total EGFR. This divergence is particularly relevant to strategies that induce LRIG1: although BMPR inhibition increased LRIG1 mRNA under hypoxia, LRIG1 protein accumulation and growth suppression were attenuated. Simply blocking lysosomal function is unlikely to solve this problem, as bafilomycin A1 also prevents LRIG1-mediated EGFR downregulation and growth suppression in colorectal cancer organoids [11]. Identifying the mechanisms underlying LRIG1 proteostasis as well as the strategies to preserve LRIG1 protein levels without impairing its ability to downregulate RTKs may therefore be necessary for LRIG1-based therapeutic approaches.

## CRediT authorship contribution statement

**Akari Minamiura:** Methodology, Investigation, Formal analysis, Visualization, Writing – original draft. **Shuto Aoki:** Methodology, Investigation, Formal analysis. **Rio Matsuwaka:** Methodology, Investigation. **Arisa Yoshimoto:** Methodology, Investigation. **Yumi Nobori:** Methodology, Investigation. **Takahiro Matsuura:** Methodology, Investigation. **Shinji Takamatsu:** Supervision. **Eiji Miyoshi:** Funding acquisition, Supervision, Writing – review & editing. **Jumpei Kondo:** Conceptualization, Funding acquisition, Methodology, Investigation, Formal analysis, Visualization, Writing – original draft.

## Ethics declaration

This study was conducted under approval from the Institutional Ethics Committee of the University of Osaka Hospital (No. 21129) and in accordance with the Declaration of Helsinki.

## Data availability

The RNA-seq data are available from the NCBI Sequence Read Archive (SRA) under BioProject accession number PRJNA1537312.

## Acknowledgement

We acknowledge the NGS core facility at the Research Institute for Microbial Diseases of the University of Osaka for the sequencing and data analysis. The authors are grateful to Prof. Masahiro Inoue (Kyoto University) for kindly providing the colorectal cancer organoids used in this study.

## Declaration of generative AI and AI-assisted technologies in the manuscript preparation process

During the preparation of this work, the authors used an AI-based language model (Perplexity, powered by GPT-5.1) to improve the language, readability, and organization of the manuscript. After using the AI tool, the authors reviewed and edited the content as needed and take full responsibility for the content of the published article.

