## Supplementary material for "Hypoxia reduces induced LRIG1 protein despite sustained mRNA induction in colorectal cancer organoids": Fig. S

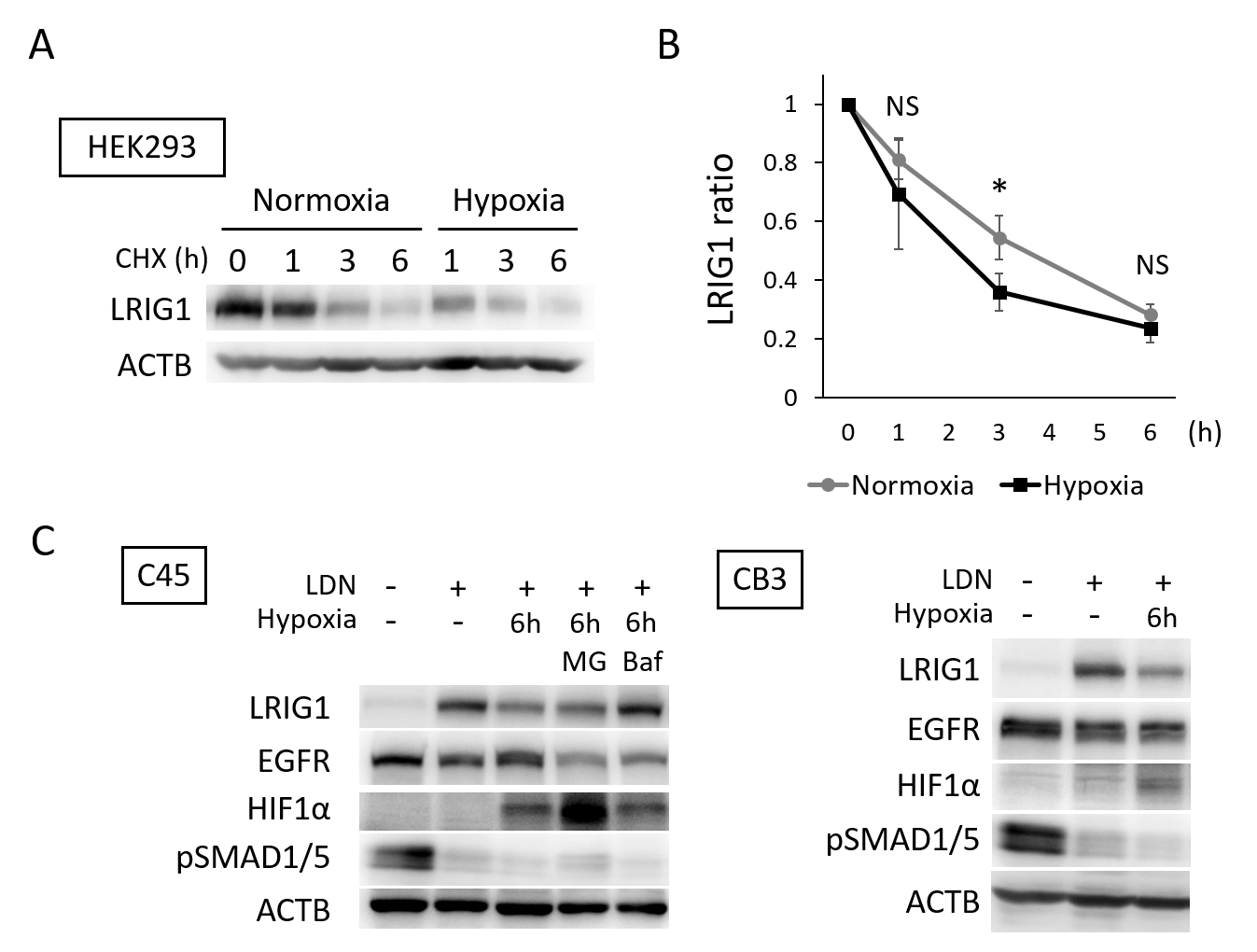

Supplementary Figure S1. Hypoxia enhances LRIG1 loss without reducing EGFR protein levels

(A-B) Cycloheximide chase assay in HEK293 cells expressing exogenous LRIG1 under normoxia or hypoxia. (A) Representative western blot. (B) Densitometric quantification from three independent experiment. *p < 0.05, NS; not significant, t-test with Bonferroni correction. (C) Western blot analysis of C45 and CB3 organoids cultured under normoxia and hypoxia with or without 10µM MG132 or 10nM bafilomycin A1.
