## Supplementary material for "Hypoxia reduces induced LRIG1 protein despite sustained mRNA induction in colorectal cancer organoids": Table S4

**Supplementary Table. S4**

|  |  |  | LRIG1 | |  | Global expression | |  | Partial correlation (LRIG1) | |
| --- | --- | --- | --- | --- | --- | --- | --- | --- | --- | --- |
|  | N |  | r | p |  | r | p |  | estimate | ｐ |
| ESCA | 181 |  | **-0.451** | <0.001 |  | -0.243 | 0.007 |  | **-0.410** | <0.001 |
| STAD | 412 |  | **-0.542** | <0.001 |  | -0.252 | <0.001 |  | **-0.509** | <0.001 |
| COADREAD | 592 |  | -0.218 | <0.001 |  | -0.042 | 0.909 |  | -0.218 | <0.001 |
| PAAD | 177 |  | **-0.484** | <0.001 |  | -0.275 | <0.001 |  | **-0.449** | <0.001 |
| LIHC | 366 |  | -0.272 | <0.001 |  | 0.338 | <0.001 |  | -0.347 | <0.001 |
| CHOL | 36 |  | -0.216 | 0.206 |  | -0.056 | 0.912 |  | -0.218 | 0.202 |
| BRCA | 1082 |  | **-0.551** | <0.001 |  | -0.146 | <0.001 |  | **-0.559** | <0.001 |
| LUAD | 510 |  | **-0.532** | <0.001 |  | -0.116 | 0.054 |  | **-0.523** | <0.001 |
| PRAD | 493 |  | -0.109 | 0.031 |  | -0.112 | 0.060 |  | -0.145 | 0.002 |
| HNSC | 515 |  | **-0.508** | <0.001 |  | **-0.436** | <0.001 |  | -0.345 | <0.001 |
| GBM | 160 |  | -0.277 | 0.001 |  | -0.059 | 0.912 |  | -0.275 | <0.001 |
| LGG | 514 |  | **-0.452** | <0.001 |  | 0.186 | <0.001 |  | **-0.431** | <0.001 |
| SKCM | 443 |  | -0.217 | <0.001 |  | -0.094 | 0.192 |  | -0.197 | <0.001 |

Table 1. Detailed data related to Figure 1B.

Correlation between hypoxia score and LRIG1 expression, correlation between global RNA expression and LRIG1 expression, and partial correlation between hypoxia score and LRIG1 expression adjusted for global RNA expression, across cancer types. Correlation coefficients (r) and partial correlation coefficient estimates (estimate) with an absolute value of 0.4 or greater are shown in bold.

N, sample number; r, correlation coefficient; p, Holm-corrected p-value; estimate, partial correlation coefficient estimate.
